## Supporting Information for "Balancing the supply and demand for taxonomy: an analysis of European taxonomic capacity and policy needs"

The following Supporting Information is available for this article:

**Figure S1.** A word cloud to visualise the most common words occurring in title and abstract of the 33,499 articles found. This gives a visual representation of the subjects of the articles and a qualitative check that we have predominantly filtered for taxonomic articles.

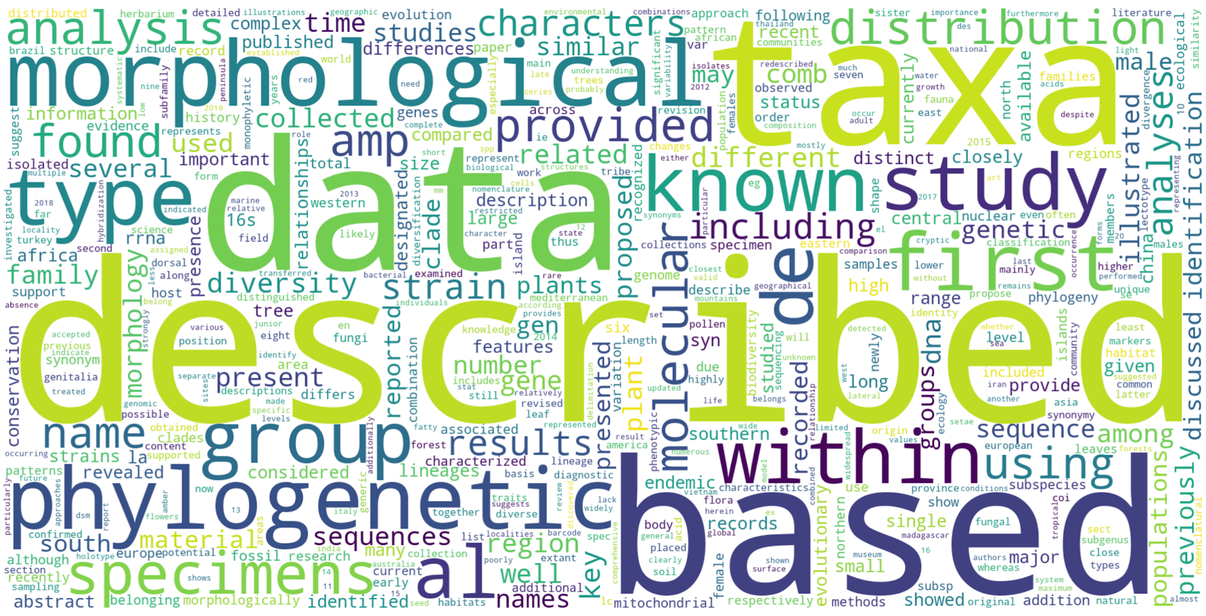


**Figure S2.** The cumulative frequency curve of newly discovered authors using 402 journals after author deduplication (see methods).

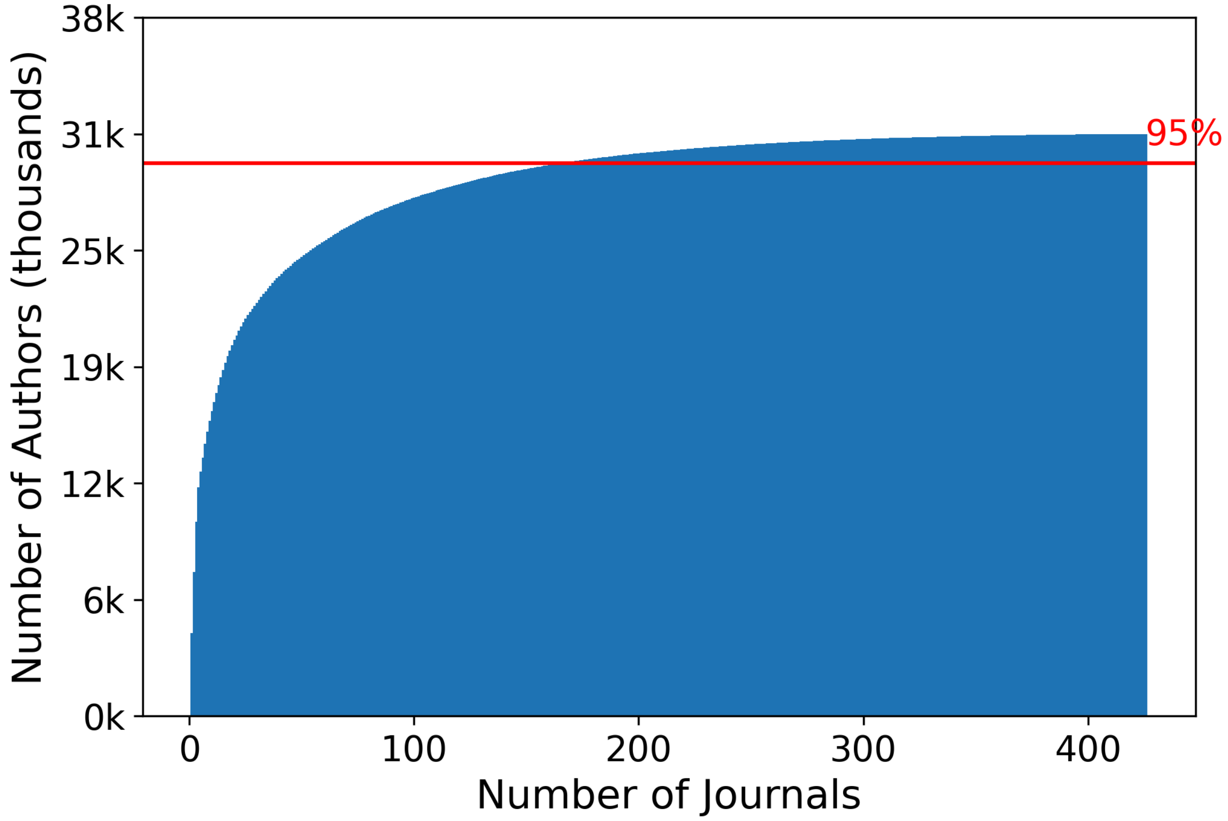


**Figure S3.** A histogram showing the familial focus of taxonomic articles with authors affiliated to European institutions. The top 10 families by number of articles are Asteraceae: 400, Staphylinidae: 311, Fabaceae: 297, Orchidaceae: 297, Poaceae: 262, Scarabaeidae: 173, Curculionidae: 168, Lamiaceae: 150, Erebidae: 145, Caryophyllaceae: 139.

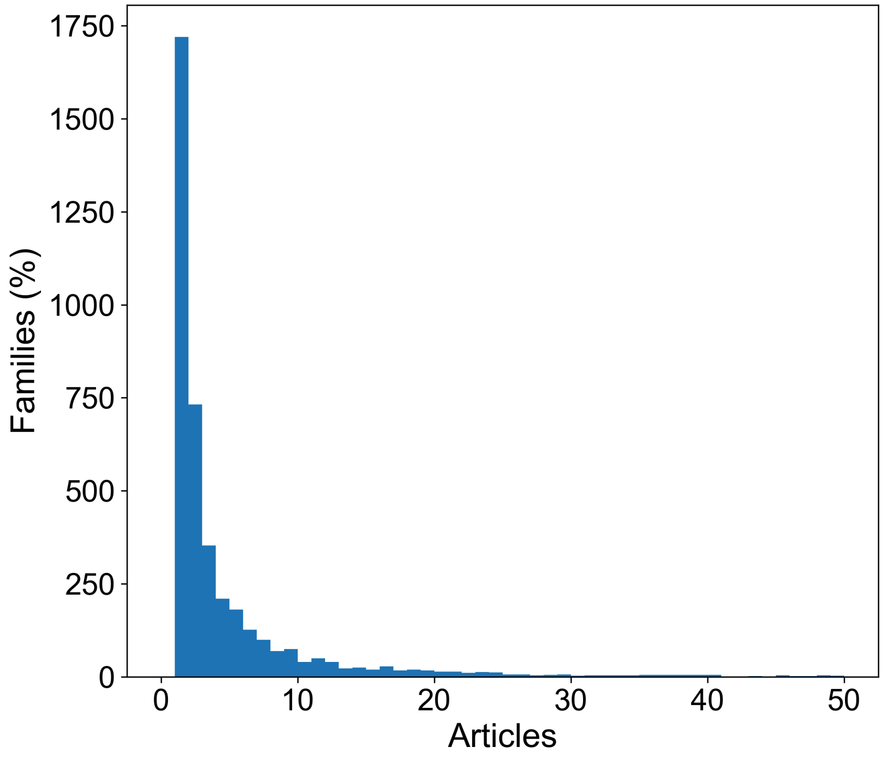


**Figure S4.** Correlation of population size and number of taxonomists per country (N = 48) in Europe. Pearson correlation coefficient: 0.8640 (p = 2.634e-15).

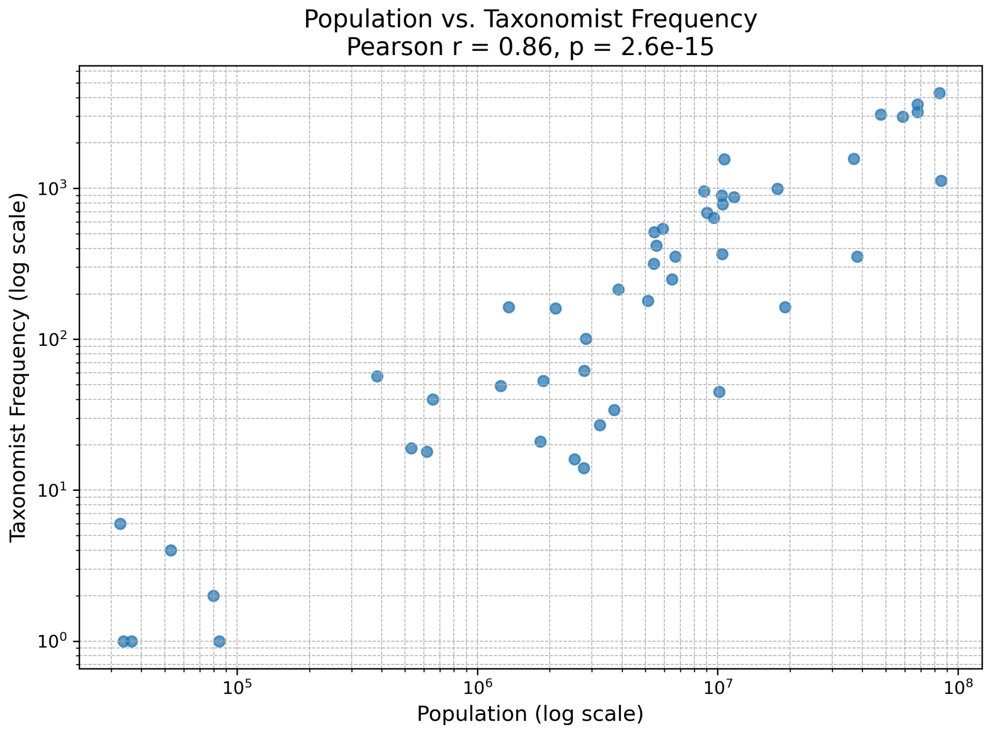


**Figure S5.** The distribution of distinct dwc:recordedby text strings (A) and dwc:identifiedBy text strings (B) on occurrences from GBIF per country from years between 2014 and 2023 inclusive. Note that the colour ramp is on a logarithmic scale and that the maps use an equal area Mollweide projection. Made with Natural Earth. Free vector and raster map data @ naturalearthdata.com.

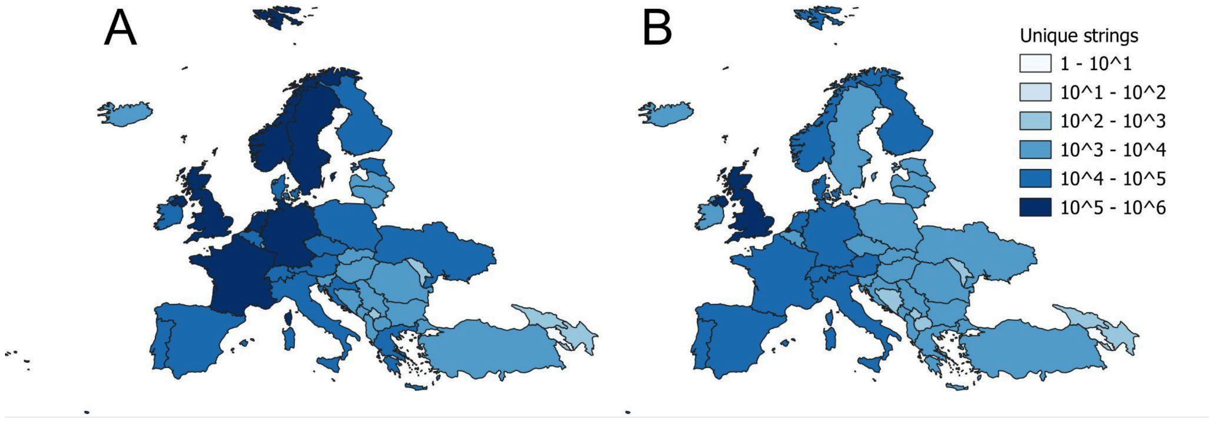

**Figure S6.** Summary diagnostics and regression results for the log–log robust regression model.

(A) Estimated regression coefficients (±95% confidence intervals) for the model relating taxonomic research effort (number of authors) to species richness and policy variables. Predictors are ordered by absolute importance.
(B) Relative importance of predictors based on the absolute value of their standardised regression coefficients.
(C) Standardised residuals from the robust regression model, with horizontal red lines indicating thresholds for potential outliers (|residual| > 3).
(D) Q–Q plot showing the quantiles of the residuals against the theoretical quantiles of a normal distribution. Deviation from the 45° reference line indicates departures from normality.

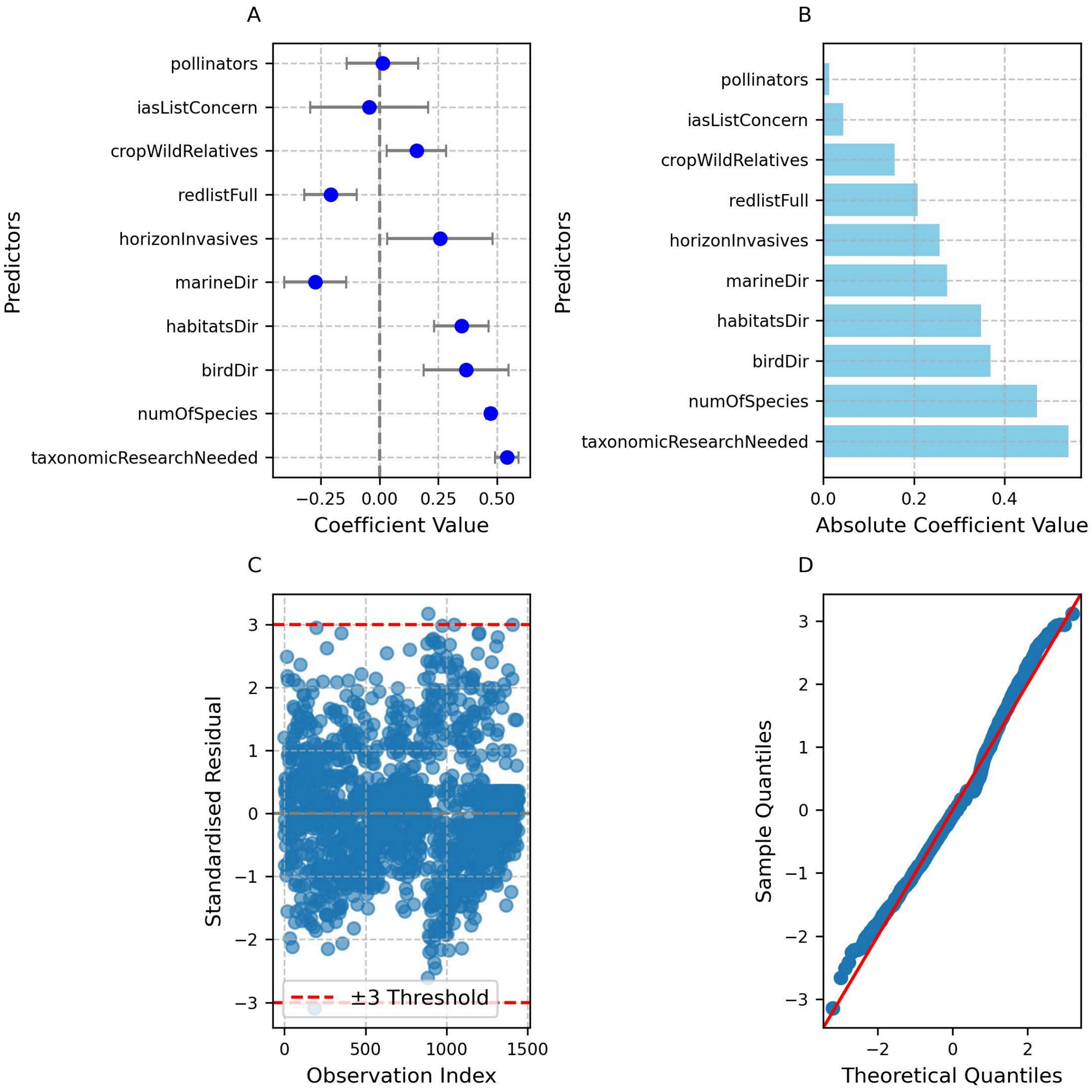

**Figure S7.** Scatterplots showing the correlation between the policy relevance of each taxonomic order and the number of taxonomists working on them, with separate panels for plants (N = 6), animals (N = 8), and fungi (N = 2). Each plot corresponds to a different policy included in this study.

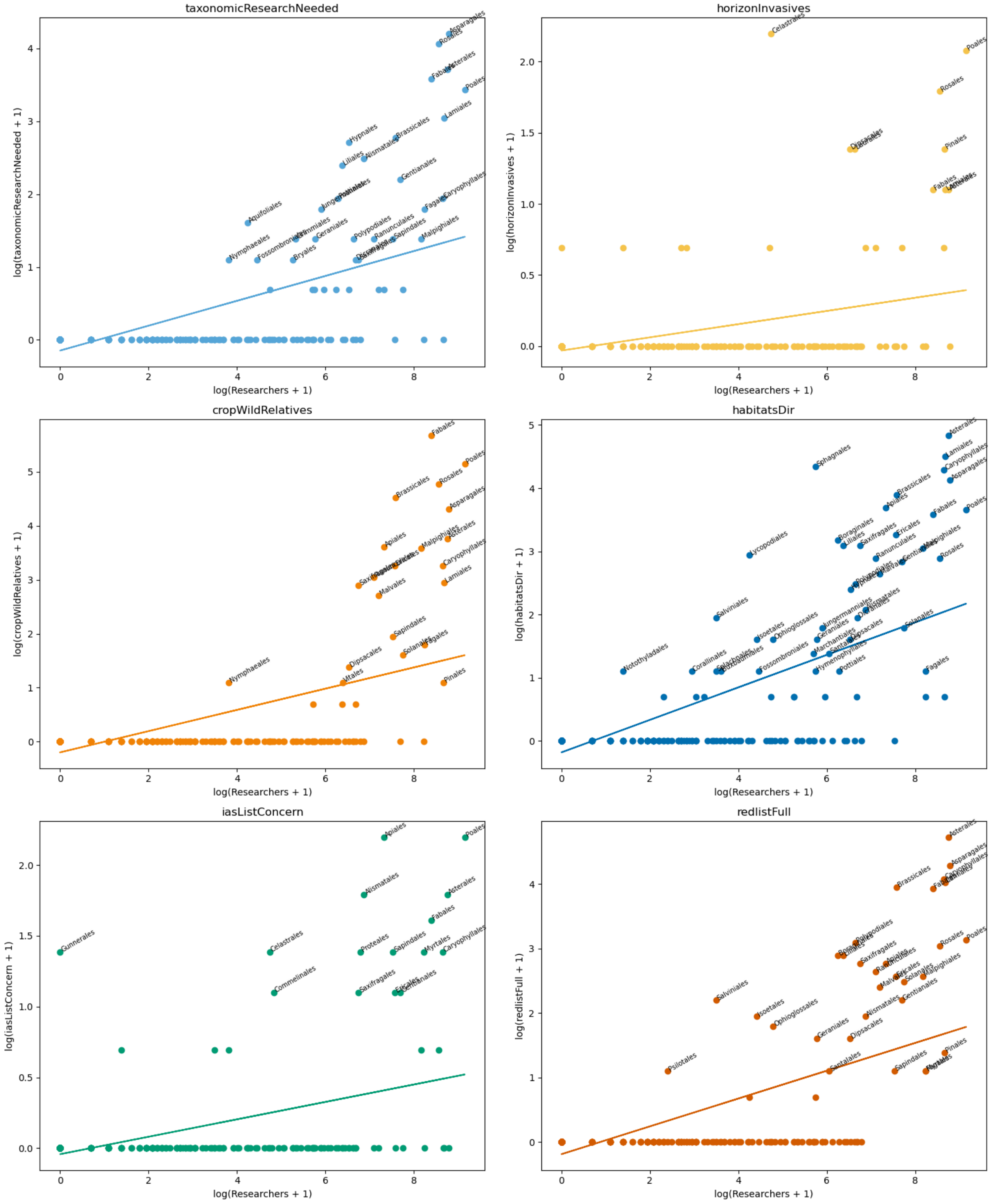

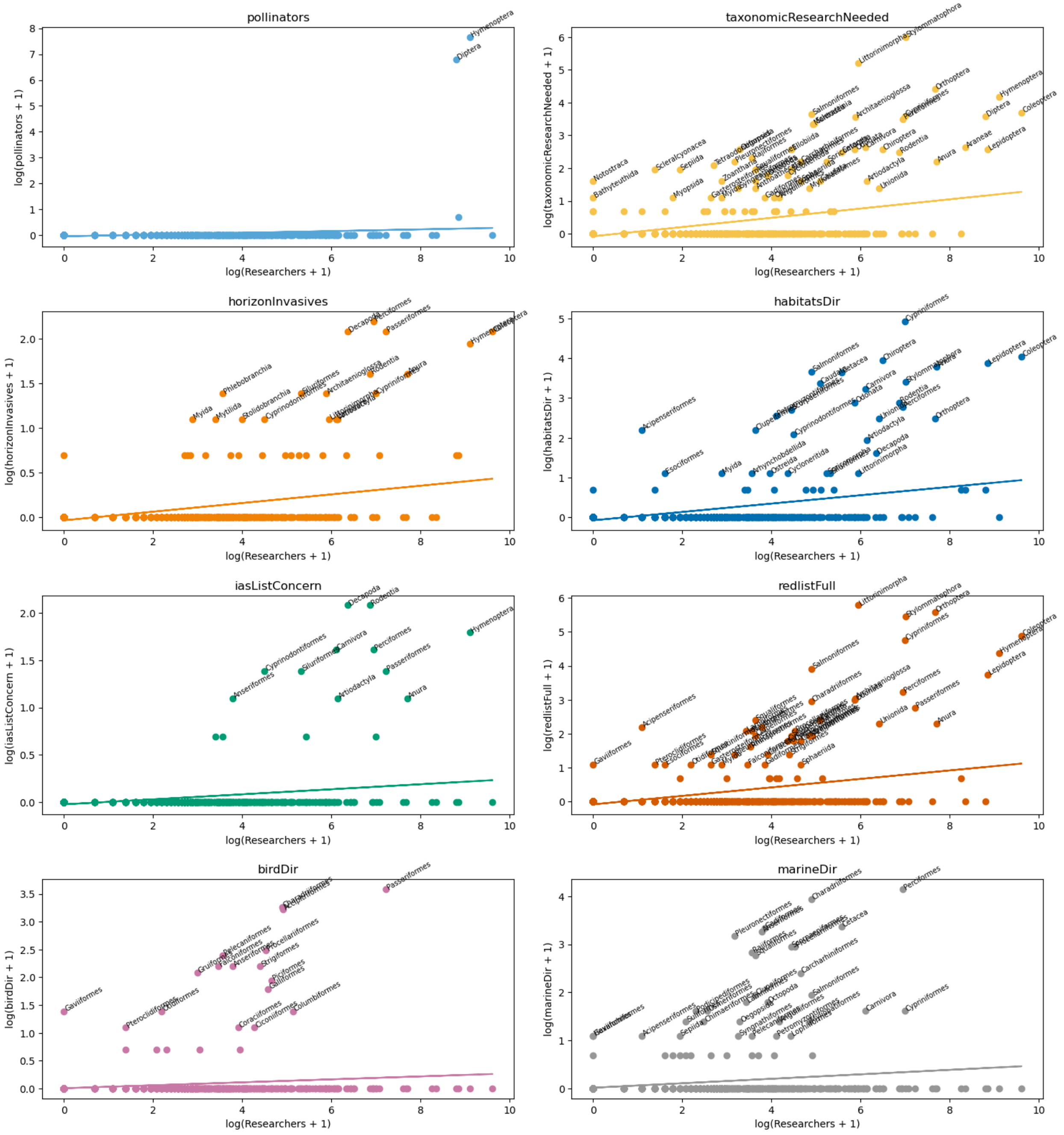
pollinators
log(pollinators + 1)
log(Researchers + 1)
taxonomicResearchNeeded
log(taxonomicResearchNeeded + 1)
log(Researchers + 1)
horizonInvasives
log(horizonInvasives + 1)
log(Researchers + 1)
habitatsDir
log(habitatsDir + 1)
log(Researchers + 1)
iasListConcern
log(iasListConcern + 1)
log(Researchers + 1)
redlistFull
log(redlistFull + 1)
log(Researchers + 1)
birdDir
log(birdDir + 1)
log(Researchers + 1)
marineDir
log(marineDir + 1)
log(Researchers + 1)

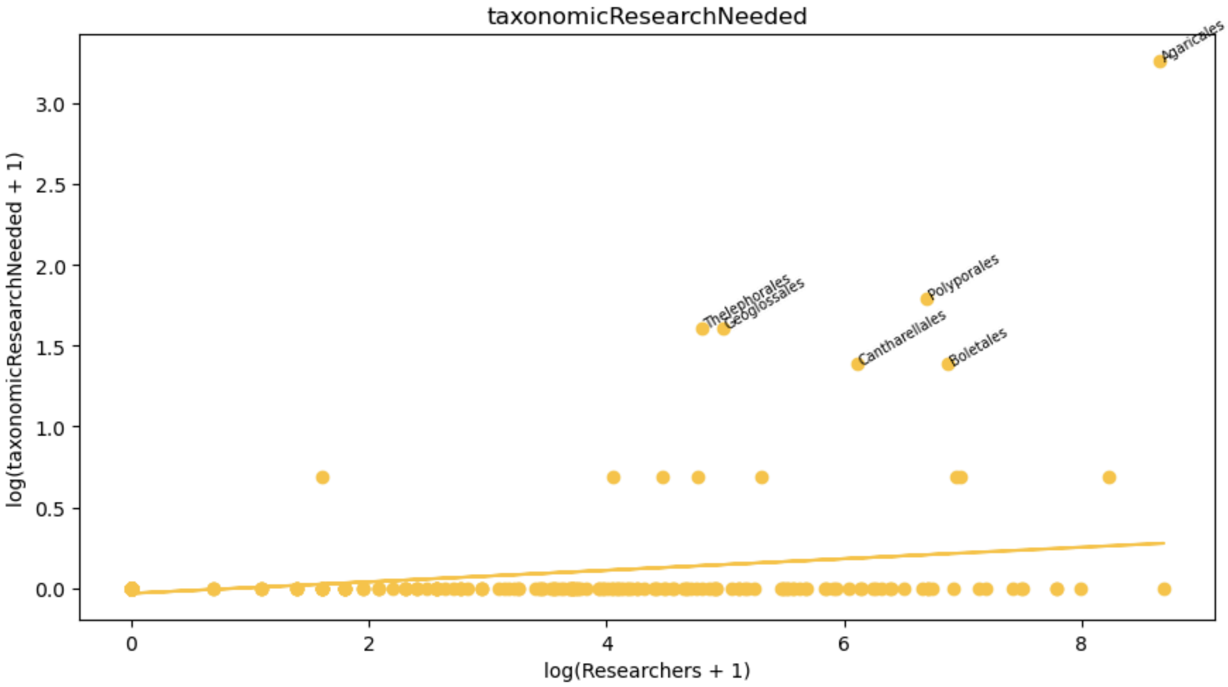
taxonomicResearchNeeded
log(taxonomicResearchNeeded + 1)
log(Researchers + 1)
Agaricales
Thelephorales
Geoglossales
Polyporales
Cantharellales
Boletales

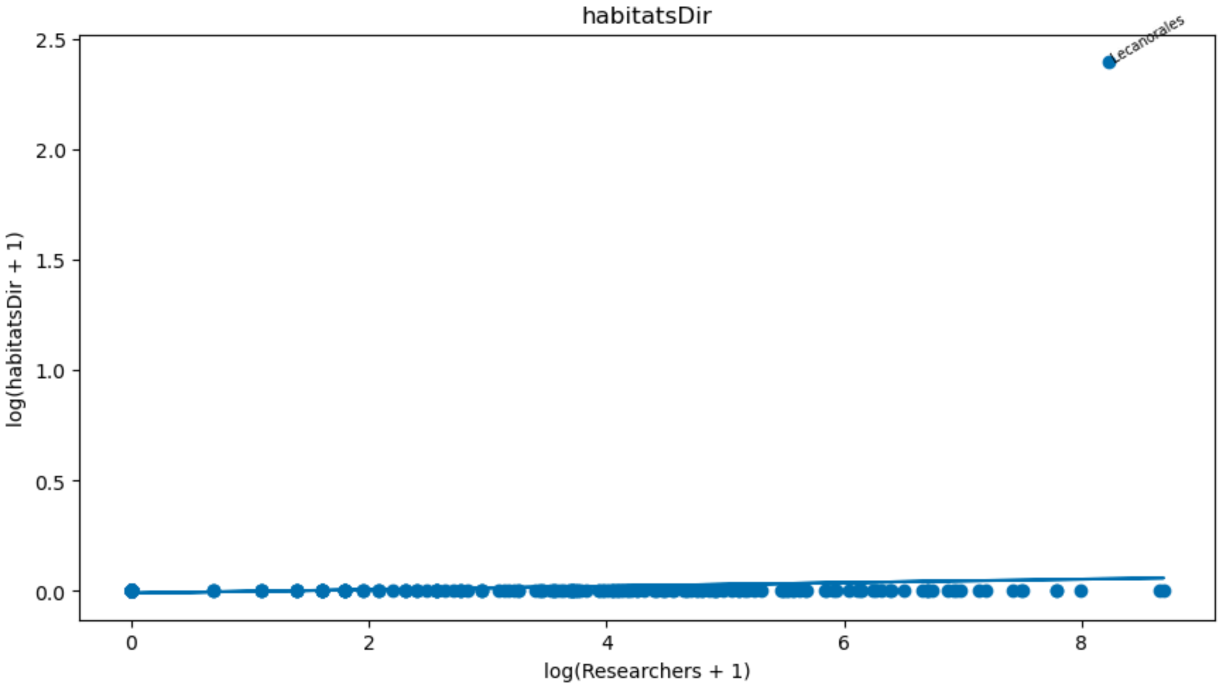
habitatsDir
log(habitatsDir + 1)
log(Researchers + 1)
Lecanorales

**Table S1:** The most common taxa in the European red list, the European red list taxonomic research needed category, the list of European crop wild relatives and invasive species on a horizon scanning list. The number of each species in each taxa are in parentheses.

| Rank | Red List - top ten families | Taxonomic research needed - top ten families | Crop wild relatives - top ten genera | Invasive species on the horizon - top nine families |
|---|---|---|---|---|
| 1 | Hydrobiidae (588) | Hydrobiidae (80) | Trifolium (71) | Formicidae (5) |
| 2 | Apidae (561) | Hydnaceae (71) | Medicago (24) | Cichlidae (4) |
| 3 | Andrenidae (465) | Moitessieriidae (55) | Vicia (21) | Poaceae (4) |
| 4 | Megachilidae (442) | Geomitridae (54) | Brassica (19) | Sciuridae (4) |
| 5 | Hygromiidae (397) | Tettigoniidae (42) | Avena (18) | Ampullariidae (3) |
| 6 | Tettigoniidae (350) | Hygromiidae (41) | Lathyrus (15) | Asteraceae (3) |
| 7 | Acrididae (334) | Amaryllidaceae (41) | Lotus (15) | Celastraceae (3) |
| 8 | Halictidae (314) | Rosaceae (40) | Prunus (15) | Cerambycidae (3) |
| 9 | Cerambycidae (278) | Clausiliidae (39) | Aegilops (14) | Fabaceae (3) |
| 10 | Cyprinidae (238) | Helicidae (37) | Allium (14) | – |
